# Human discrimination and categorization of emotions in voices: a functional Near-Infrared Spectroscopy (fNIRS) study

**DOI:** 10.1101/526558

**Authors:** T. Gruber, C. Debracque, L. Ceravolo, K. Igloi, B. Marin Bosch, S. Frühholz, D. Grandjean

## Abstract

Variations of the vocal tone of the voice during speech production, known as prosody, provide information about the emotional state of the speaker. In recent years, functional imaging has suggested a role of both right and left inferior frontal cortices in attentive decoding and cognitive evaluation of emotional cues in human vocalizations. Here, we investigated the suitability of functional Near-Infrared Spectroscopy (fNIRS) to study frontal lateralization of human emotion vocalization processing during explicit and implicit categorization and discrimination. Participants listened to speech-like but semantically meaningless words spoken in a neutral, angry or fearful tone and had to categorize or discriminate them based on their emotional or linguistic content. Behaviorally, participants were faster to discriminate than to categorize and they processed the linguistic content of stimuli faster than their emotional content, while an interaction between condition (emotion/word) and task (discrimination/categorization) influenced accuracy. At the brain level, we found a four-way interaction in the fNIRS signal between condition, task, emotion and channel, highlighting the involvement of the right hemisphere to process fear stimuli, and of both hemispheres to treat anger stimuli. Our results show that fNIRS is suitable to study vocal emotion evaluation in humans, fostering its application to study emotional appraisal.

---

Much of the initial vocal emotional processing in the brain occurs in subcortical and sensory cortical areas for a review, see ^1,2^. However, many higher order processes occur in cortical areas, including the associative temporal and the prefrontal cortices ^3–7^. For example, in recent years, a large implication of the prefrontal cortex (PFC) has been suggested for the processing of emotional stimuli in the vocal and auditory domain, based on work conducted mainly with functional Magnetic Resonance Imaging (fMRI) ^8–10^. In particular, the inferior frontal gyrus (IFG) is involved in the processing of human vocal sounds, and reacts to some of its properties such as prosody, the variation in intonations that modulates vocal production ^3,11^. Connectivity analyses such as psycho-physiological interactions have revealed that the bilateral IFGs are functionally connected to sensory and secondary temporal cortical areas ^8,12^. The right IFG appears especially activated during the listening of emotional stimuli ^5^. In comparison, activations of the left IFG have been connected to the semantic content of a given vocal utterance, in part because the left IFG encompasses Broca’s area, which is particularly involved in speech processing ^13^. Nevertheless, this lateralized view of the activity of the IFG is not shown in all studies. Indeed, several studies on emotional processing have found bilateral activations of the IFG ^3,14,15^, or even left activations of specific areas of the IFG ^5,16,17^ during emotional tasks. This suggests that different areas of the two IFGs are involved in different tasks concerned with the treatment of emotional vocal stimuli ^18^. In a recent meta-analysis, Belyk et al. ^4^ have reviewed the role of the *pars orbitalis* of the IFG during semantic and emotional processing, highlighting a possible functional organization in two different zones. The lateral one, close to the Broca’s area, would be involved in both semantic and emotional aspects while the ventral frontal operculum would be more involved in emotional processing per se. The lateral zone would have been co-opted in human communication for semantic aspects while in non-human primates this zone would be more related to emotional communication. While we broadly agree with this view, the potential existence of vocalizations with semantic content in non-human primates ^19,20^ suggests that this co-optation may have emerged earlier in our evolution.

One specific domain where the IFG appears to be involved is the categorization and discrimination of emotions in auditory stimuli, both implicitly and explicitly. Implicit processing occurs when participants are required to conduct a task (e.g. judging the linguistic content of words or sentence pronounced with different emotional tones) other than evaluating the emotional content of the stimuli e.g. ^3,17^. The IFG is also involved when participants make explicit judgments (e.g. categorizing anger vs fear) about the emotional content of the stimuli they are exposed to ^12,21–23^. The right IFG may be particularly important for conducting such an explicit evaluation of the emotional content of the voices, although both hemispheres play a role in the processing of the emotional content ^18^. In particular, experiments often make use of non-words or pseudo-words, that is utterances that have the linguistic structure of words but without their semantic content (e.g. ‘belam’ or ‘molem’), such that the linguistic structure of these stimuli can trigger activity in the left hemisphere, both during explicit and implicit judgment tasks ^13^.

While the majority of the studies investigating cognitive processes in cortical regions have relied on fMRI or electroencephalography (EEG), studies utilizing functional Near-Infrared Spectroscopy (fNIRS) have developed over the last twenty-five years ^24–30^. fNIRS has recently also proven a useful non-invasive technique to study emotion processes ^31^, especially in the visual domain for a review, see ^32^. For example, fNIRS has been used to study affective processing of pictures in the parietal and occipital areas ^33^ and recent work suggests that a large occipital-parietal-temporal network is involved in discrimination tasks involving judgments about ‘emotional’ gait patterns ^34^. So far, PFC activations have been recorded during two types of task: the passive viewing and active categorization of emotional stimuli. In the first case, researchers found an increase of the oxygenated hemoglobin (Oxy-Hb) in the bilateral ventrolateral PFC when participants were watching negative pictures; in contrast, positive pictures led to a decrease of Oxy-Hb in the left dorsolateral PFC ^35^. In the second case, the authors isolated an activation of the bilateral PFC involving an increase of Oxy-Hb and a decrease of deoxygenated hemoglobin (Deoxy-Hb) when participants were viewing fearful rather than neutral images ^36^. These results are consistent with recent findings showing fNIRS activations in ventrolateral PFC during the viewing of threatening pictures ^37^. However, some studies did not find differences in Oxy-Hb between baseline and any kind of pictures, whether negative, neutral or positive ^38^. A natural negative mood during task completion was also found to have an impact on PFC activity during a working memory task ^39^, although an experimentally induced negative mood had the opposite effect with increased PFC Oxy-Hb ^40^. As of now, the emerging picture for affective visual stimuli is that the PFC is solicited during both passive and active stimulation; however, the exact pattern of activity must be characterized with more studies and with an effort towards more comparability between the paradigms employed across fNIRS studies ^32^.

In the present study, our goal was to investigate whether fNIRS could be used to measure blood oxygen-level dependent (BOLD) signal variations after implicit and explicit judgments of the emotional content of vocal utterances. One of the major advantages of fNIRS over fMRI is the fact that it is noiseless. This is all the more important in the acoustic domain where fMRI results will always be affected by scanner noise effects. In this study we set out to study differences in BOLD signal over the IFG in relation to emotional voices in a perfectly silent environment. While much attention has been devoted to the study of visual emotional stimuli using fNIRS as described above, to our knowledge, only two studies have been published on the treatment of vocal emotional stimuli and both showed that emotional stimuli activated the auditory cortex more compared to neutral stimuli ^41,42^. While Plichta and colleagues did not investigate how vocal emotional stimuli modulated the activity in the PFC, Zhang and colleagues showed that the left IFG was modulated by emotional valence (positive vs negative) and they also found a bilateral activation for the orbito-frontal cortex (OFC) when anger was contrasted with neutral stimuli (42). However, none of these two studies investigated categorization and discrimination of vocal emotional stimuli, making unclear whether fNIRS could be used for such analysis. Our goal here was thus to assess whether fNIRS was suitable to study the discrimination and categorization of auditory stimuli at the level of the IFG ^43^. To do so, we applied paradigms developed in fMRI studies relying on the use of pseudo-words and requiring participants to perform explicit judgment tasks either on the linguistic or emotional content of negative or neutral vocal stimuli. We first predicted that active tasks (categorization and discrimination) would activate the IFG differently compared to passive listening of auditory stimuli. Second, we predicted that the fNIRS signal would be modulated differentially with the experimental manipulation of the task (categorization or discrimination) and the content focus (condition: word or emotion). More specifically we predicted that categorization (processing A-versus-B computations) would lead to a larger recorded brain activity because it is cognitively more demanding than discrimination (only processing A-versus-Non-A computations). Finally, we investigated the predicted differential IFG lateralization related to linguistic and emotion processing.

## Material and Methods

### Participants

Twenty-eight healthy volunteers (14 males; mean age 26.44 years, SD=4.7, age range 21-35) took part in the experiment. The participants reported normal hearing abilities and normal or corrected-to-normal vision. No participant presented a neurological or psychiatric history, or a hearing impairment. All participants gave informed and written consent for their participation in accordance with the ethical and data security guidelines of the University of Geneva. The study was approved by the Ethics Cantonal Commission for Research of the Canton of Geneva, Switzerland (CCER).

### Stimuli

The stimulus material consisted of three speech-like but semantically meaningless two-syllable words (i.e. “minad”, “lagod”, “namil”) ^44^. These pseudowords were 16-bit recordings sampled at a 44.1 kHz sampling rate, presented at 70 dB Sound Pressure Level (SPL). Two male and two female speakers spoke these three different pseudowords in an angry, fearful, or neutral tone, resulting in a total of 36 individual stimuli. From a pre-evaluation of all stimuli on five emotion scales (sadness, joy, anger, fear, neutral), we selected three pseudowords for each emotional tone that were most consistently evaluated as angry, fearful, and neutral, respectively ^44^ and see Appendix.

### Procedure

Participants seating in front of a computer performed two alternative forced-choice tasks of auditory discrimination (A vs non-A) and categorization (A vs B) via keyboard pressing. Stimuli were presented binaurally through in-ear headphones (Sennheiser). The participants listened to each voice and made a corresponding button press as soon as they could identify the requested target for each block. The categorization and discrimination blocks were split into blocks with a focus on emotion and blocks with a focus on the linguistic word features of the stimuli. Our experiment was thus blocked by tasks, based on a two (task: discrimination/categorization) by two (condition: emotion/word) design, with two blocks per condition and task (two each for emotion categorization, word categorization, emotion discrimination, and word discrimination). These eight blocks were preceded and followed by passive listening blocks, leading to ten blocks in total (Figure 1). During the two passive blocks, participants only had to listen to the stimuli without having to make an active decision. Button assignments, target button and target stimuli were counterbalanced across blocks for each participant. Task blocks also alternated across the experiment, and block order and response buttons were counterbalanced across participants.

The two blocks of emotion categorizations involved a three-alternative forced-choice determining whether the speaker’s voice expressed an “angry”, “fearful”, or “neutral” tone (the options “angry” and “fear” were assigned to left and right index finger button, the “neutral” option included a simultaneous press of the left and right button no more than 500ms apart).

The two blocks of word categorization involved a three-alternative forced-choice determining whether the word spoken was “minad”, “lagod”, or “namil” (the options “minad” and “lagod” were assigned to left and right index finger button, the “namil” option included a simultaneous press of the left and right button no more than 500ms apart).

The discrimination blocks included a target emotion or a target word, which was assigned to one of the two response buttons. During the two emotion discrimination blocks, either angry or fearful voices were the target (e.g. press the left button for “angry” voices, and the right button for all other voices) and the two word discrimination blocks included either “minad” or “lagod” as the target word (e.g. press the left button for “minad”, and the right button for all other words).

Within each block, all 36 voice stimuli were presented twice resulting in 72 trials per block. These 72 trials were clustered into mini-blocks of six voice stimuli, where a stimulus was presented every 2s; each mini-block thus had an average length of 11.5-12s. The presentation of mini-blocks was separated by 10s blank gap for BOLD signal to return to baseline. Trials for each mini-blocks were randomly assigned, with the only exception that every emotion (with no more than three times the same emotion in a row) and every word had to appear at least one time per mini-block. Each mini-block started with a visual fixation cross (1×1°) presented on a grey background for 900±100ms. The fixation cross prompted the participant’s attention and remained on the screen for the duration of the mini-block.

**Figure 1.**
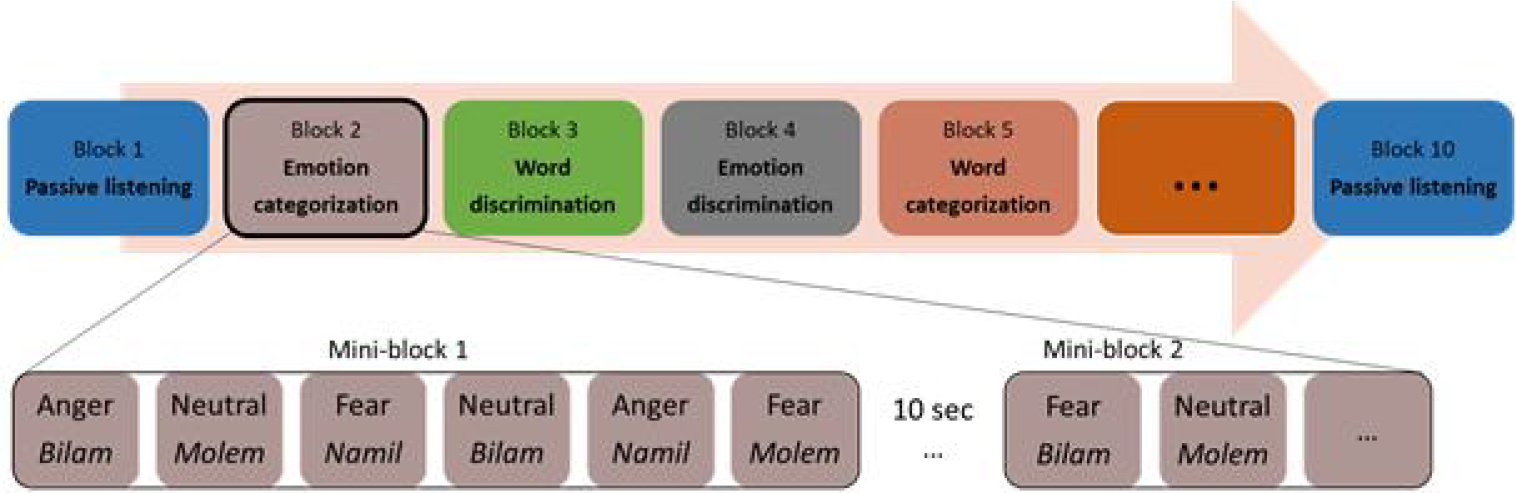
Experimental protocol with a possible list of blocks and stimuli within a mini-block.

### NIRS recordings

fNIRS is as a non-invasive and non-ionizing method to study the brain hemodynamics ^28^. Using the principle of tissue transillumination, fNIRS measures via near infrared lights blood oxygenation changes related to the hemodynamic response function (HRF) constituted of Oxy-Hb and Deoxy-Hb. For this study, we used the Oxymon MKIII device (Artinis Medical Systems B.V., Elst, The Netherlands) with a 2×4 optode template and wavelengths of 765 nm and 855 nm. We placed four optodes as a square on both sides of the participant’s head, forming 4 channels around the F7 or F8 references and corresponding respectively to the left and right inferior frontal gyri (Figure 2), as defined in the 10-20-electroencephalogram (EEG) system ^45,46^. All channels were placed at an inter-optode distance of 35 mm and we recorded with a sampling rate of 250Hz.

**Figure 2.**
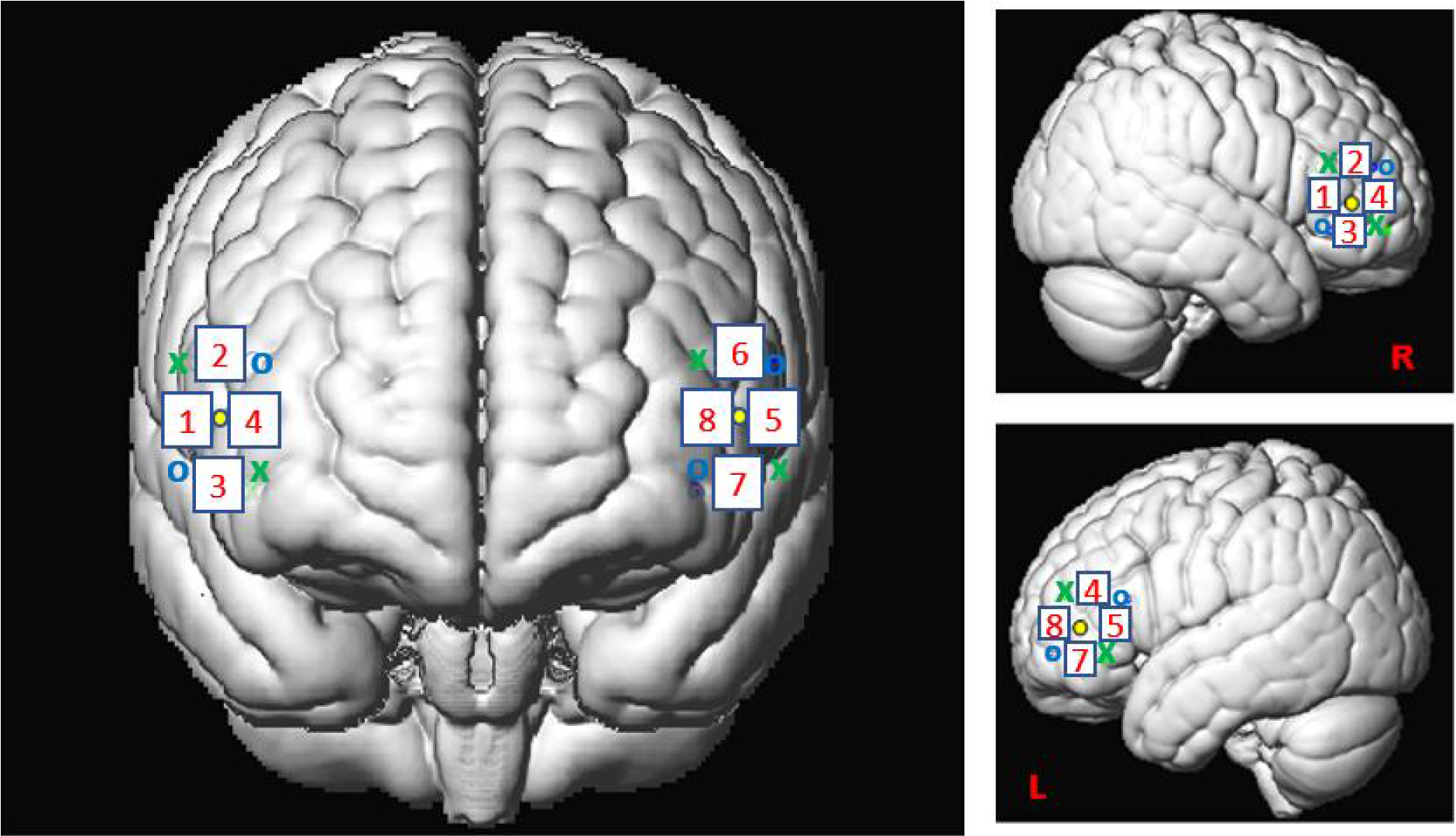
Spatial registration of optode locations to the Montreal Neurological Institute (MNI) space using spatial registration approach ^43^. This method relies on structural information from an anatomical database to estimate the fNIRS probe locations into a 3D space. Thus, this procedure allows the projection of the 8 channels in the subject space into the MNI ^46^. Central dots indicate the F7 and F8 electrode position in the 10-20 EEG system. “o” and “x” indicate optical source and detector positions respectively.

### Analysis

#### Behavioral data

We only analyzed data from participants who completed all the blocks (N=26, 2 excluded) using R studio software (R Studio team (2015) Inc., Boston, MA, url:http://www.rstudio.com/). We assessed accuracy in the tasks by predicting a Generalized Linear Mixed Model (GLMM) with binomial error distribution, with condition (emotion vs word), task (categorization vs discrimination) as well as their interaction as fixed factors, and with participant IDs and blocks (first or second) as random factors, against a GLMM with the same factors but not including the interaction between condition and task. This analysis was followed by contrasts for which post-hoc correction for multiple comparisons was applied by using a Bonferroni correction (0.05/4=0.0125). We analyzed reaction times by predicting a General Linear Mixed (GLM) model with condition (emotion vs word), task (categorization vs discrimination) as well as their interaction as fixed factors, and with participant IDs and blocks as random factors, against the same model without the interaction using R studio software. For the reaction time analysis, we only included cases where the participants were correct in their response.

#### fNIRS data

Seven participants out of 28 were excluded from the dataset due to poor signal quality or missing fNIRS data. A total of 21 participants were thus analyzed in this study. We performed the first level analysis with MATLAB 2016B (Mathwortks, Natick, MA) using the SPM_fNIRS toolbox (Tak, 2016; https://www.nitrc.org/projects/spm_fnirs/) and homemade scripts. Hemoglobin conversion and temporal preprocessing of Oxy-Hb and Deoxy-Hb were made using the following procedure:

i. hemoglobin concentration changes were calculated with the modified Beer-Lambert law ^47^;
ii. motion artifacts were reduced using the method proposed by Scholkmann et al. ^48^ based on moving standard deviation and spline interpolation;
iii. physiological and high frequency noise was removed using a band-stop filter between 0.12-0.35 Hz and 0.7-1.5 Hz following Oostenveld et al. ^49^ and a low-pass filter based on the HRF ^50^;
iv. fNIRS data were down-sampled to 10 Hz;
v. low frequency confound were reduced using a high-pass filter based on a discrete cosine transform set with a cut-off frequency of 1/64Hz ^50^.

We performed the second level analysis with R studio using Linear Mixed Models analysis with condition (emotion vs word), task (categorization vs discrimination vs passive) and channel (1 to 8) as well as their interactions as fixed factors, with participant IDs and block orders as random factors. In particular, we predicted models including a higher-level interaction against models of the lower dimension (e.g. a four-way versus a three-way interaction + the main effects), presented in the results, on which we ran subsequent contrasts. Models with lower dimension interactions can be found in Appendix.

#### Analyses including passive blocks

We first aimed to isolate whether our regions of interest (ROIs) were activated differently during active blocks compared to passive blocks. To do so, our first analyses confronted data collected during the passive and the active blocks. We were particularly interested in testing the effects of lateralization and emotional content, as previous fMRI studies had shown possible variation for these factors (see above). We noticed post-hoc that subjects’ activations during the first and the final passive run differed widely, with the activation pattern found during the final passive run close to the pattern of activation recorded during the active tasks (see Appendix, in particular Figure 7). Therefore, it is likely that subjects were still engaged, consciously or not, in the discrimination or categorization of stimuli during the final passive block, even though they were instructed not to do so. For this reason, we excluded data from the final passive block, and only included data from the first passive block, for which no instruction besides listening to stimuli had been conveyed to the participants, ensuring their naivety to the task. To isolate any effect of active processes (that is processes occurring during blocks where the task was either discrimination or categorization) vs passive processes, we tested a three-way model including data from the first passive run and all discrimination and categorization blocks. We specifically tested effects of active vs passive blocks across emotions and channels, resulting in testing a three-way interaction between process (active vs passive tasks), emotion (anger vs fear vs neutral) and channels (C1 to C8). Subsequently, we were interested in contrasting hemisphere rather than channel activities, that is we studied lateralization by pulling together data from channels 1-4 for the right hemisphere and data from channels 5-8 for the left hemisphere.

#### Analyses on active blocks only

Second, we were interested in whether there were differences in activations between categorization or discriminations of words and emotions across hemispheres, and whether this depended on the emotion being tested. To do so, we focused on active blocks (discrimination and categorization blocks) and excluded the passive blocks, as the subjects had no specific instructions regarding the stimuli compared to the active blocks (see above). To isolate any differences between the factors, we tested a four-way interaction on the active blocks including the effects of channels (C1 to C8), tasks (discrimination vs categorization), conditions (word vs emotion), and emotions (anger vs fear vs neutral). Subsequently, as in our first analysis, we tested contrasts between right and left hemispheres, pulling together data from C1-C4 and C5-C8 channels respectively. In a final analysis, we individually looked at each hemisphere to contrast anger, respectively fear, versus neutral stimuli.

For both analyses, we averaged the concentration of Oxy-Hb over the period of 4 to 12 s post stimulus onset for each trial. This period was selected to include the range of maximum concentration changes (μM) observed across participants for Oxy-Hb and Deoxy-Hb. We performed the same analyses on Deoxy-Hb to check our Oxy-Hb concentration changes (μM) for consistency. Because our results with Deoxy-Hb were similar to the Oxy-Hb we only provide our results for Oxy-Hb in the main text (see Appendix for Deoxy-Hb analyses). All data were log-transformed to normalize them for the analyses.

## Results

### Behavioral data

#### Accuracy data

The models included 14’200 individual data points across 26 participants. There was no significant effect for the two main factors condition (emotion vs word; χ^2^(1)= 0.62, p=0.43) and task (categorization vs discrimination; (χ^2^(1)=0.92, p=0.34) but there was a significant interaction between condition and task (χ^2^(1)=18.68, p<0.001). Following the Bonferroni correction, the following contrasts were significant: emotion/word categorization (χ^2^(1)=12.24, p<0.001); emotion/word discrimination (χ^2^(1)=6.65, p<0.01); emotion categorization/discrimination (χ^2^(1)=13.21, p<0.001); and there was a tendency for word categorization/discrimination (χ^2^(1)=5.87, p=0.015). On average, participants were more accurate when engaging in emotion categorization (97.91% accuracy) than word categorization (96.58% accuracy), but more accurate when engaged in word discrimination (97.50%) than emotion discrimination (96.46%, Figure 3).

**Figure 3.**
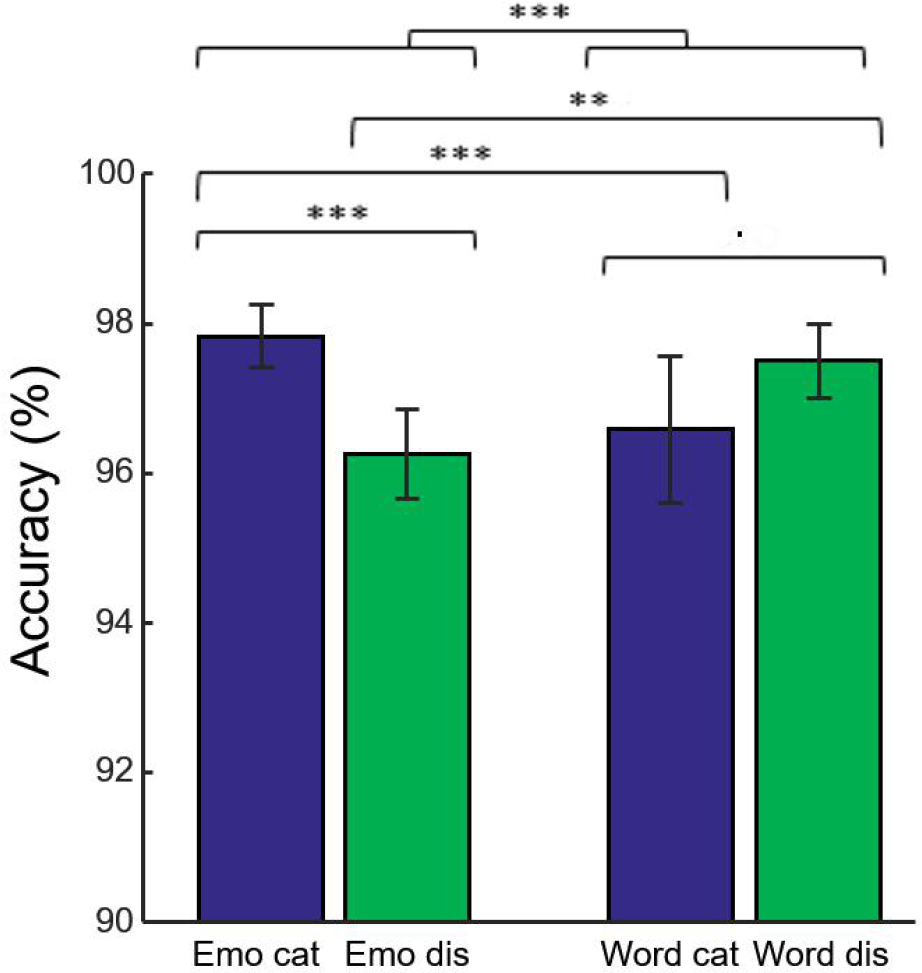
Accuracy (in %) of the 26 participants represented as a function of condition (emotion vs word) and task (categorization vs discrimination).

#### Reaction time

Correlation between reaction time and accuracy was extremely weak (Spearman’s rho=0.033). All the analyses below were only done on correct trials; the models included 13’789 individual data points across 26 participants. We found no interaction between the two factors (χ^2^(1)=2.61, p>0.10) but significant main effects for condition (χ^2^(1)=234.30, p<0.001) and task (χ^2^(1)=617.28, p<0.001). The mean reaction time for categorization was 1012.77ms (N=6’860), significantly longer than the mean reaction time for discrimination (897.10ms, N=6’929). Similarly, average reaction time was 987.34ms (N=6’873) for emotion content, which was significantly longer than 922.16ms (N=6’916) for the word modality (Figure 4).

**Figure 4.**
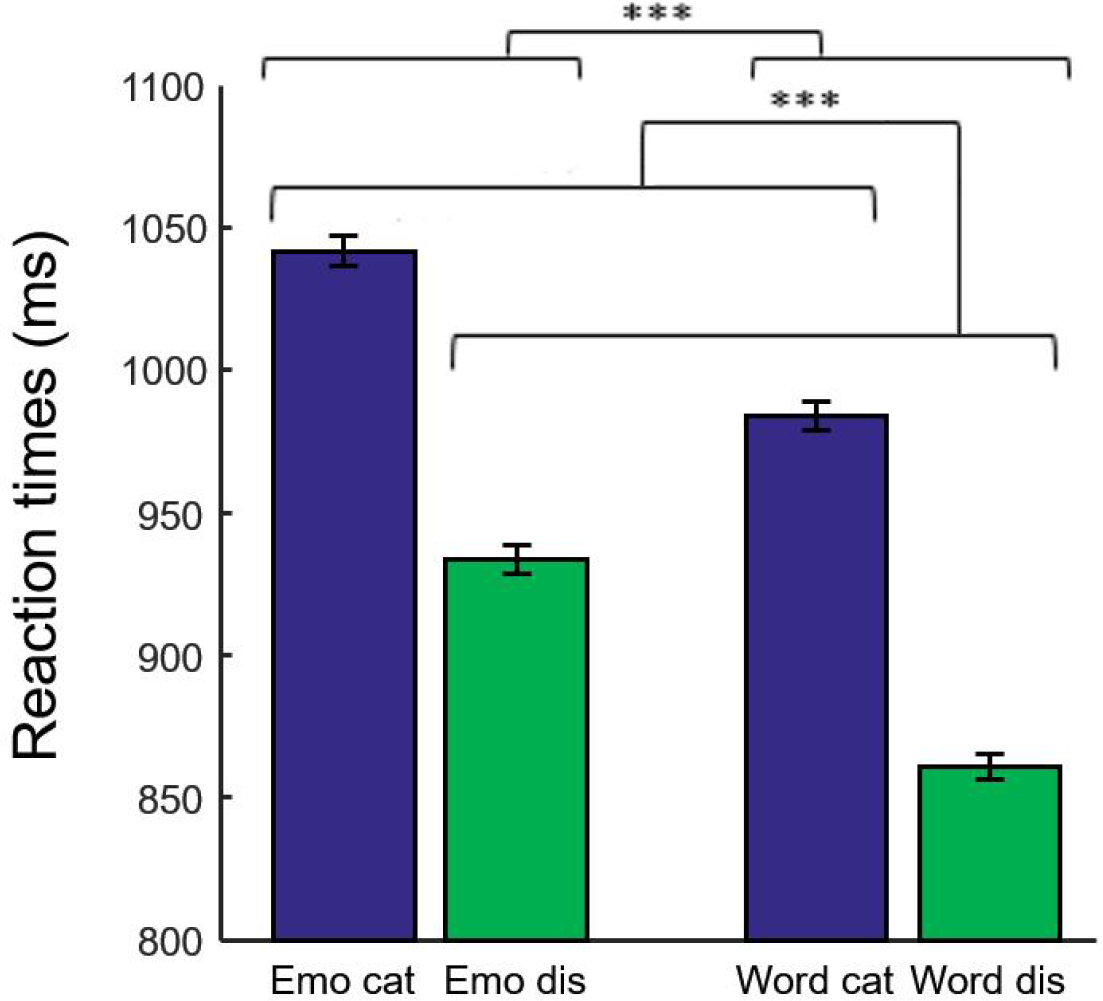
Reaction time (in ms) for the correct trials of the 26 participants represented as a function of condition (emotion vs word) and task (categorization vs discrimination).

### NIRS data

#### Analyses including the first passive run

As predicted, we revealed a significant three-way interaction of task by channel by emotion (χ^2^(46)=7246.20, p<0.001). This three-way interaction was followed-up by the following contrasts: while the contrast passive vs tasks (categorization and discrimination) with lateralization (left, C5–C6–C7–C8 vs right, C1–C2–C3–C4) and ‘anger’ vs ‘neutral’ did not yield significant differences (χ^2^(1)=0.14, p=0.71), there was a significant difference with higher Oxy-Hb values for tasks vs passive listening for ‘fear’ compared to ‘neutral’ on the right compared to left hemisphere (χ^2^(1)=18.51, p<0.001; Figure 5). The contrast between passive listening and tasks was also significant with higher values for left compared to right when only neutral stimuli were considered (χ^2^(1)=29.62, p<0.001; see Figure 8 in Appendix).

**Figure 5.**
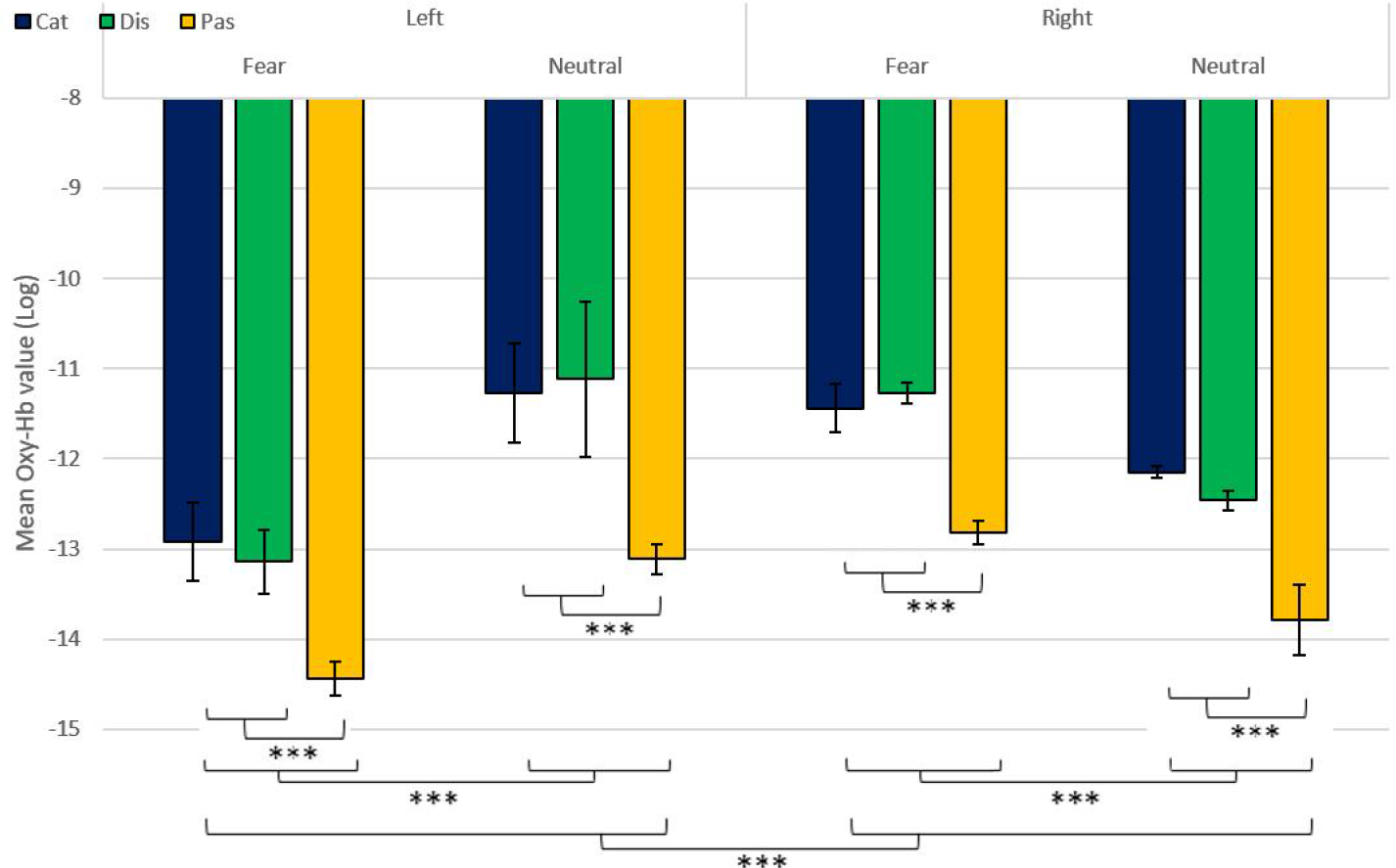
Contrast in log of Oxy-Hb concentration changes (μM) in the right and left hemispheres during the treatment of fear and neutral stimuli.

#### Analyses of the active blocks

We revealed a significant four-way interaction of task by channel by condition by emotion (χ^2^(62)=7204.3, p<0.001), confirmed also for Deoxy-Hb (χ^2^(62)=3117.5, p<0.001, see Appendix). To test the specific significant effects related to emotions and lateralization, we performed the following contrasts: we tested the impact of condition (emotion versus word), lateralization (left vs right) and task (discrimination vs categorization) for anger (χ^2^(1)=2.79, p=0.095), fear (χ^2^(1)=11.75, p<0.001), and neutral (χ^2^(1)<1). The contrasts condition, lateralization, task for anger vs fear (χ^2^(1)=1.55, p=0.21) and anger vs neutral (χ^2^(1)=1.81, p=0.18) did not reach significance but the contrast fear vs neutral did (χ^2^(1)=6.69, p<0.01, Figure 6).

**Figure 6.**
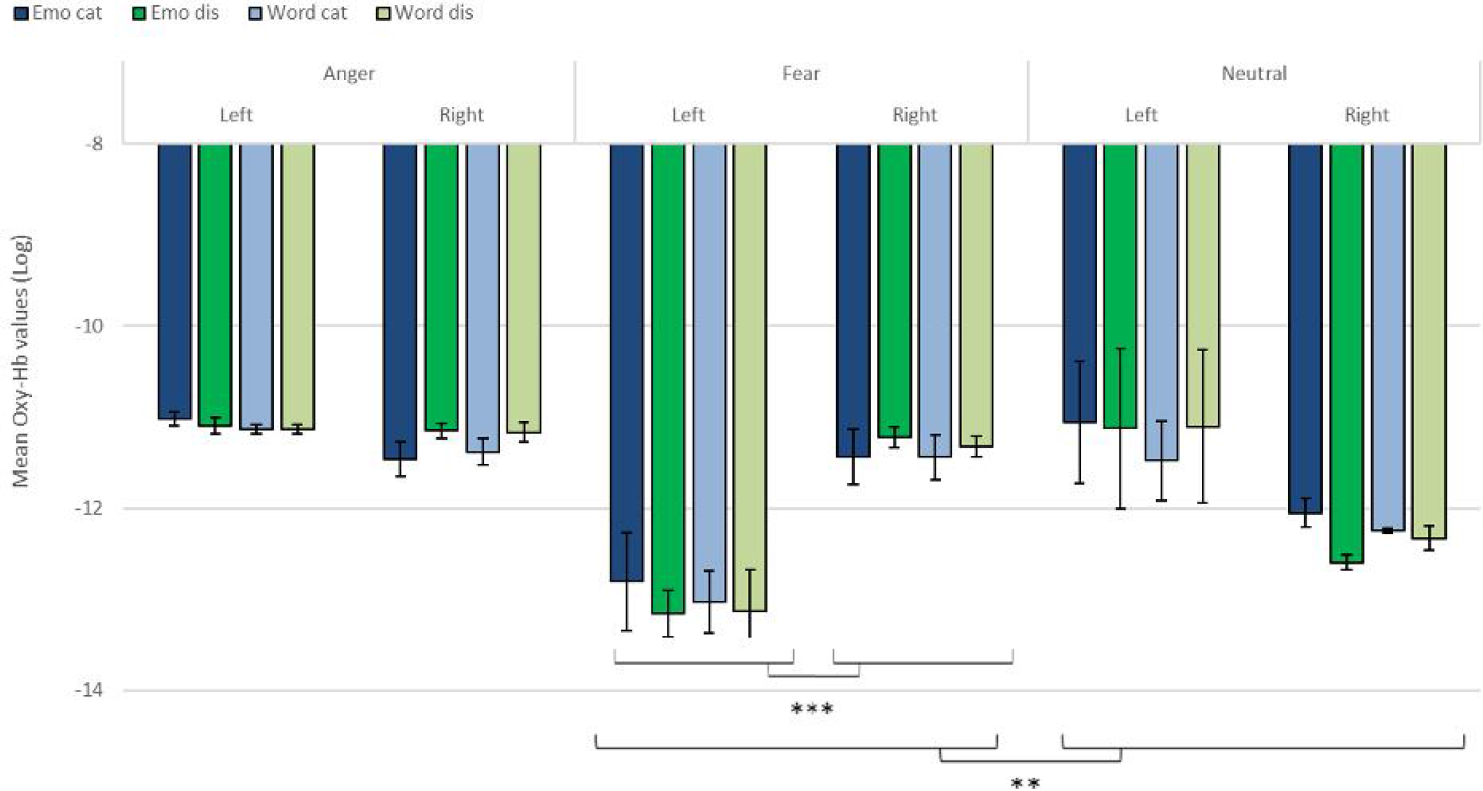
Contrast in log of Oxy-Hb concentration changes (μM) for anger, fear and neutral stimuli in the right and left hemispheres for emotional categorization/discrimination and word categorization/discrimination.

To investigate the specificities of the lateralization, we also ran contrasts on the left or right channels only. This analysis revealed a significant effect on the left channels for ‘anger’ vs ‘neutral’ (χ^2^(1)=12.32, p<0.001); the effect was also significant for the right channels (χ^2^(1)=29.28, p<0.001), with both effects going towards the same direction. The comparison for ‘fear’ vs ‘neutral’ when only the left channels were considered was marginal, possibly due to the differences of contingency of variances (χ^2^(1)=3.04, p=0.081) while its was significant for the right channels (χ^2^(1)=29.18, p<0.001).

## Discussion

In this study we showed that fNIRS is a suitable method to study cognitive paradigms related to emotions, particularly categorization and discrimination, in the human prefrontal regions. Our first goal was to estimate whether it was possible to isolate significant activity in the IFG, whose activity has been highlighted in previous fMRI studies investigating emotional prosody processing, and in particular during categorization and discrimination of emotional stimuli ^3,11^. Both right and left IFGs have been connected to the processing of emotional stimuli ^3,5,51^ and we were interested in replicating and extending such effects using fNIRS. While fNIRS cannot specifically target a small region such as IFG because of its limited spatial accuracy, we positioned our channels so that any variation in BOLD signal would be most parsimoniously attributed to variation of activity in this area.

We found significant differences in frontal activations during our tasks, including a difference in activation during categorization and discrimination compared to passive listening in the Oxy-Hb and confirmed in the Deoxy-Hb signals. In particular, in our first analysis of the NIRS signal, we isolated left hemisphere activity for active processing versus passive listening of neutral stimuli. This result suggests that fNIRS is a generally suitable method to identify signatures of categorization and discrimination in auditory stimuli, independently of their emotional content. Furthermore, the results of our second analysis revealed significant differences between categorization and discrimination between the experimental conditions. This is exemplified with the significant four-way interaction between condition (word vs emotion), task (categorization vs discrimination), emotion and channel. Interestingly, the results of the contrasts suggest that differences in brain activity between categorization and discrimination and lateralization are more important for fear stimuli compared to neutral ones. Conversely, the lack of overall significance for the contrast between hemispheres for anger versus neutral stimuli appears to result from the fact that effects went into the same direction in the two hemispheres, as shown when angry and neutral stimuli were compared independently in our final analysis. In addition, activity for anger stimuli across conditions and tasks was higher compared to other stimuli (less negative in Figure 6). This result supports a bilateral approach to the treatment of emotional stimuli ^7,11^. In contrast, the effect for fear versus neutral was more pronounced in the right hemisphere, in line with classic studies highlighting a right dominance for emotional treatment in prosody ^5^.

Our behavioral results are also informative with respect to a differential treatment of stimuli depending on condition and task. While our participants were generally accurate across tasks and conditions (over 96% correct in all tasks), we found minor but significant variations between the four experimental conditions, which may reflect the variations in treatment outlined in the four-way interaction found in the fNIRS data. Participants were most accurate when engaged in emotional categorization, seconded by word discrimination, with the lowest accuracy rates found for word categorization and emotional discrimination. This result may seem counter-intuitive at first, as categorization appears to be cognitively more difficult than discrimination. However, participants’ reaction times also varied between the conditions: overall, categorization took more time compared to discrimination, with judgements made on emotional content always taking longer than on linguistic content. Combined, these results suggest different processing between words and emotions (in line with ref. 4), with active judgements on emotional stimuli being more demanding (longest reaction time) than judgements on the linguistic content. Indeed, when participants judged the emotional content of stimuli, they were more accurate for categorization than discrimination but spent a longer time before selecting their answer. In contrast, for words, participants were more accurate for discrimination compared to categorization, but they spent less time before answering.

Another potential explanation for the differences observed between the active processing of emotional aspects compared to linguistic aspects lies in the fact that the IFG is activated during both implicit and explicit categorization and discrimination of emotions ^3,10,12,17,21,22^. Our participants may thus have engaged in implicit emotional processing of the stimuli even when their task was to judge the linguistic aspect of the stimuli. This additional treatment may explain the Oxy-Hb differences found between emotions even in the context of word categorization and discrimination. The right IFG has previously been highlighted as particularly important in the explicit evaluation of the emotional content of the voices, and our Oxy-Hb results support this view, particularly when considering fear versus neutral stimuli. The generally higher activity in both hemispheres when participants processed stimuli with an angry content also supports the view that both hemispheres play a role in the processing of the emotional content, whether implicit or explicit ^18^. Future work will need to explore the specific aspects of emotional stimuli when more types of emotion (e.g. positive) are included. It may also be interesting to study whether bilateral or unilateral treatments are elicited depending of the evaluation process, implicit or explicit.

To conclude, our study shows that fNIRS is a suitable method to study emotional auditory processing in human adults with no history of psychiatric antecedents or hearing impairment. Beyond fNIRS studies investigating emotions from a perceptual point of view e.g. ^41,42^, our study replicates effects found with more traditional imaging methods such as fMRI and shows that subtle differences can be found in fNIRS signal across tasks and modalities in the study of emotional categorization and discrimination. Future work will need to examine in more details whether differences between stimuli valence or arousal may also influence the fNIRS signal. Additionally, the portability and easiness of use of fNIRS may help extend such results in populations where the use of fMRI is limited such as young infants, populations in less developed countries or, possibly, other species ^20^. The use of unfamiliar non-verbal human or nonhuman vocalizations rather than pseudo-words may be particularly informative to study the developmental and evolutionary origins of the two cognitive processes. fNIRS may also be very informative in the context of prosody production thanks to its resistance to movement artifacts compared to other brain imaging methods. Finally, our study contributes to the growing field of affective neurosciences, confirming through a different imaging technique that emotion treatment, both explicitly and implicitly, may be largely conducted in the IFG, a possible hub for the extraction and detection of variant/invariant aspects of stimuli (e.g. acoustical features) subjected to categorization/discrimination representation (e.g. anger/neutral prosody) in the brain.

## Acknowledgments

We thank the SNSF for supporting the National Center of Competence in Affective Sciences (NCCR Grant 51NF40-104897 to DG) hosted by the Swiss Center for Affective Sciences.

## Appendix

## Selection of stimuli for the experiment

From a pre-evaluation of all stimuli on five emotion scales (sadness, joy, anger, fear, neutral), we selected three pseudowords for each emotional tone that were most consistently evaluated as angry (*F*_4,80_=256.111, *p*<0.001), fearful (*F*_4,80_=151.894, *p*<0.001), and neutral (*F*_4,80_=193.527, *p*<0.001), respectively ^44^. One-way ANOVA revealed that there was a statistically significant difference in arousal scores among the angry, fear, and neutral voices (*F*_2,40_=54.073, *p*<0.001). Bonferroni corrected post-hoc planned comparisons revealed that arousal scores for angry and fearful tones were significantly higher than scores received by neutral tones (*p*<0.001), but they did not differ significantly from each other (*p*<=0.408).

## Analyses first (active and passive) blocks versus second (active and passive) blocks

Task * Channel
We revealed a significant interaction of task * block number (χ^2^(2)=2388.50, p<0.001).
Contrasts
Passive * active: (χ^2^(1)=2400.60, p<0.001)
Passive 1 * Passive 2: (χ^2^(1)=4.334.10, p<0.001)

**Figure 7.**
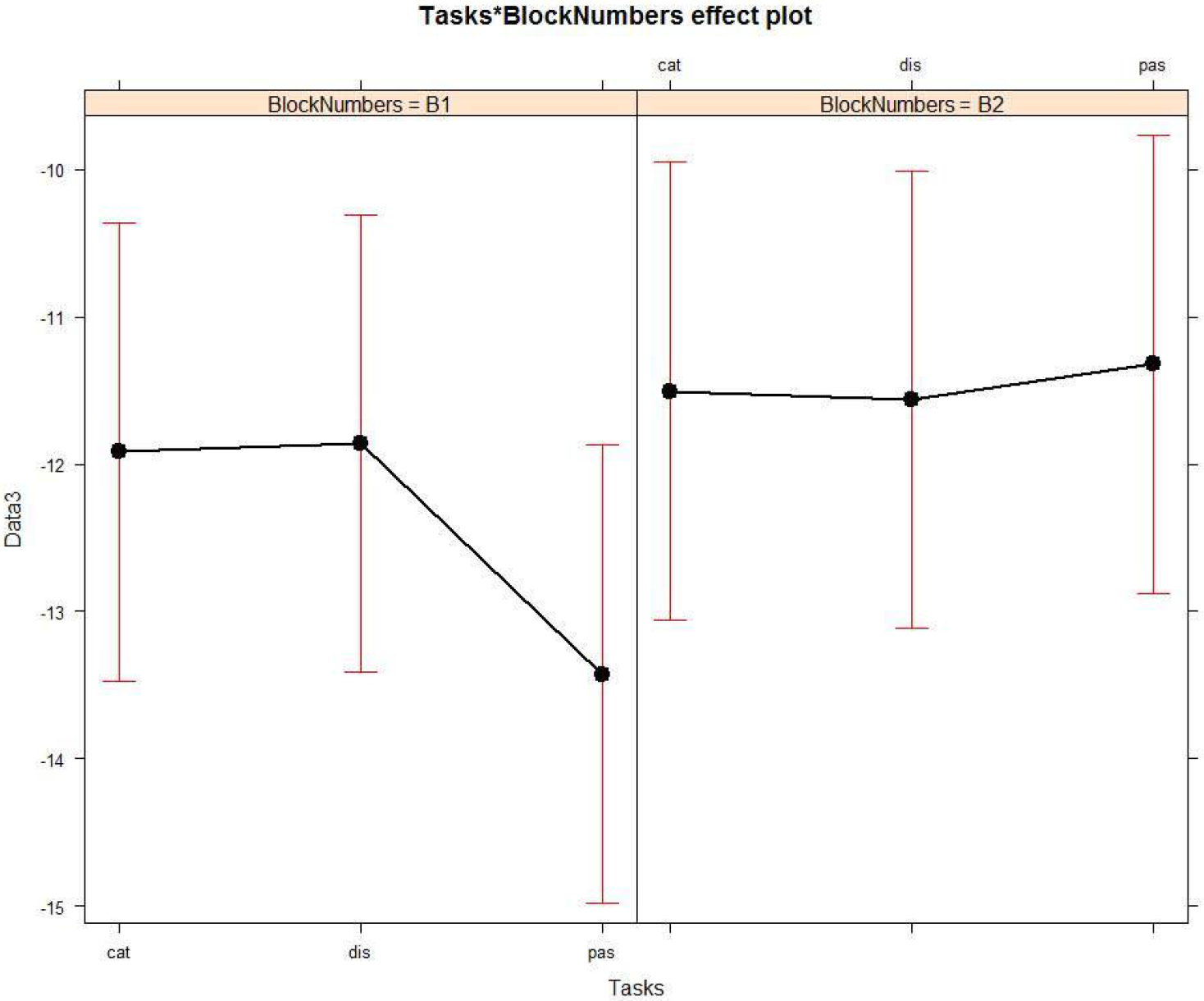
Contrast between log values of Oxy-Hb concentration changes (μM) for activities during the first (B1) and the second (B2) passive listening and active blocks including all stimuli.

## Analyses Oxy-Hemoglobin Signal

## Analyses including the first passive run

## Main effects

Channel
We revealed a significant main effect of channel (χ^2^(7)=877.25, p<0.001).
Emotion
We revealed a significant main effect of emotion (χ^2^(2)=2876.4, p<0.001).
Task
We revealed a significant main effect of task (χ^2^(2)=3617.2, p<0.001).

## Interactions

Task * Emotion
We revealed a significant two-way interaction of task * emotion (χ^2^(4)=49.21, p<0.001).
Task * Channel
We revealed a significant two-way interaction of task * channel (χ^2^(14)=177.42, p<0.001).
Emotion * Channel
We revealed a significant two-way interaction of task * channel (χ^2^(14)=6800.5, p<0.001).
Task * Channel * Emotion
We revealed a significant three-way interaction of task * channel * emotion (χ^2^(46)=7246.2, p<0.001).
Contrasts
Active vs passive, anger vs neutral, left vs right (laterality): (χ^2^(1)=0.14, p=0.71)
Active vs passive, fear vs neutral, left vs right: (χ^2^(1)=18.51, p<0.001)
Active vs passive, only neutral, left vs right: (χ^2^(1)=29.62, p<0.001)

## Analyses on active blocks

## Main effects

Condition
We revealed a significant main effect of condition (χ^2^(1)=14.84, p<0.001).
Emotion
We revealed a significant main effect of emotion (χ^2^(2)=2708.8, p<0.001).
Channel
We revealed a significant main effect of channel (χ^2^(7)=921.23, p<0.001).
Task
There was no significant effect of task (χ^2^(1)=0.01, p=0.92).

## Interactions

Task * Channel
We revealed a significant two-way interaction of task * channel (χ^2^(7)=78.001, p<0.001).
Task * Condition
We revealed a significant two-way interaction of of task * condition (χ^2^(1)=29.7, p<.001).
Emotion * Channel
We revealed a significant two-way interaction of task * emotion (χ^2^(14)=6821.9, p<0.001).
Channel * Condition
We revealed a significant two-way interaction of channel * condition (χ^2^(7)=26.31, p<0.001).
Task * Channel * Condition
We revealed a significant three-way interaction of task * channel * condition (χ^2^(15)=77.50, p<0.001).
Task * Channel * Emotion
We revealed a significant three-way interaction of task * condition * emotion (χ^2^(30)=7092.6, p<0.001).
Task * Channel * Condition * Emotion
We revealed a significant four-way interaction of task * channel * condition * emotion (χ^2^(62)=7204.3, p<0.001).
Contrasts
Task * laterality: (χ^2^(1)=5.95, p=0.015)
Task * laterality * condition: (χ^2^(1)=7.89, p<0.01)
Task * laterality * condition (anger only): (χ^2^(1)=2.79, p=0.10)
Task * laterality * condition (fear only): (χ^2^(1)=11.75, p<0.001)
Task * laterality * condition (neutral only): (χ^2^(1)=0.05, p=0.82)
Task * laterality * condition (anger vs fear): (χ^2^(1)=1.55, p=0.21)
Task * laterality * condition (anger vs neutral): (χ^2^(1)=1.81, p=0.18)
Task * laterality * condition (fear vs neutral): (χ^2^(1)=6.69, p<0.01)
Task * channels C1-C4 (right) * condition (anger vs neutral): (χ^2^(1)=29.28, p<0.001)
Task * channels C5-C8 (left) * condition (anger vs neutral): (χ^2^(1)=12.32, p<0.001)
Task * channels C1-C4 (right) * condition (fear vs neutral): (χ^2^(1)=29.18, p<0.001)
Task * channels C5-C8 (left) * condition (fear vs neutral): (χ^2^(1)=3.04, p=0.081)
Task * channels C1-C4 (right) * condition (anger vs fear): (χ^2^(1)<1, p=.993)
Task * channels C5-C8 (left) * condition (anger vs fear): (χ^2^(1)=3.12, p=.077)

## Analyses Deoxy-Hemoglobin Signal

## Analyses including the first passive run

## Main effects

Channel
We revealed a significant main effect of channel (χ^2^(7)=477.5, p<0.001).
Emotion
We revealed a significant main effect of emotion (χ^2^(2)=2632.1, p<0.001).
Task
We revealed a significant main effect of task (χ^2^(2)=1807.7, p<0.001).

## Interactions

Task * Emotion
We revealed a significant two-way interaction of task * emotion (χ^2^(4)=153.63, p<0.001).
Task * Channel
We revealed a significant two-way interaction of task * channel (χ^2^(14)=77.21, p<0.001).
Emotion * Channel
We revealed a significant two-way interaction of emotion * channel (χ^2^(14)=2890.8, p<0.001).
Task * Emotion * Channel
We revealed a significant three-way interaction of task * emotion * channel (χ^2^(46)=3293.5, p<0.001).

## Analyses on active blocks

## Main effects

Condition
We revealed a significant main effect of condition (χ^2^(1)=24.06, p<0.001).
Emotion
We revealed a significant main effect of emotion (χ^2^(2)=2628.3, p<0.001).
Channel
We revealed a significant main effect of channel (χ^2^(7)=444.14, p<0.001).
Task
We revealed a significant main effect of task (χ^2^(1)=83.42, p<0.001).

## Interactions

Task * Channel
We revealed a significant two-way interaction of task * channel (χ^2^(7)=47.08, p<0.001).
Task * Condition
There was no significant effect of task * condition (χ^2^(1)=1.62, p=0.20).
Task * Emotion
We revealed a significant two-way interaction of task * emotion (χ^2^(2)=76.56, p<0.001).
Channel * Condition
We revealed a significant two-way interaction of channel * condition (χ^2^(7)= 18.86, p=0.0086) Task *channel * Emotion
We revealed a significant three-way interaction of task * channel * emotion (χ^2^(30)=3029.5, p<0.001).
Task * Channel * Condition
We revealed a significant three-way interaction of task * channel * condition (χ^2^(15)= 32.06, p=0063).
Task * Channel * Condition * Emotion
We revealed a significant four-way interaction of task * channel * condition * emotion (χ^2^(62)=3117.5, p<0.001).

**Figure 8.**
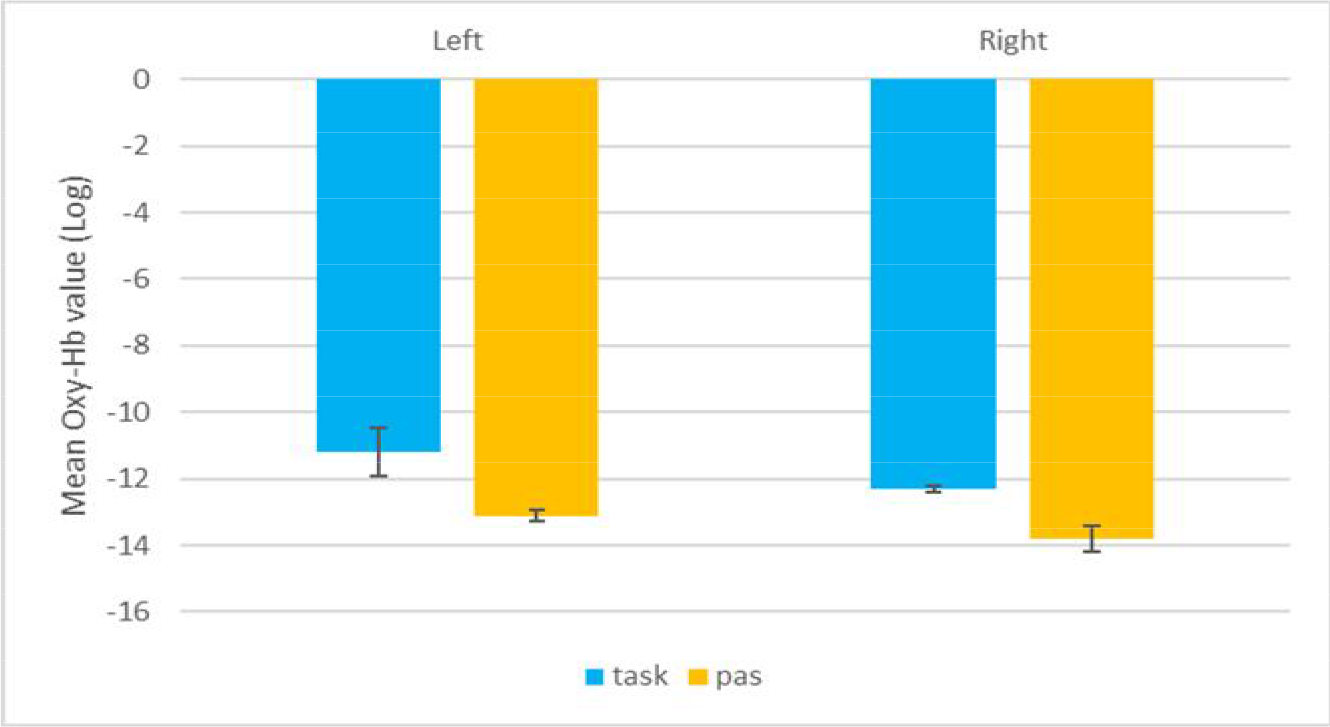
Contrast between log values of Oxy-Hb concentration changes (μM) for activities during passive listening and active (categorisation and discrimination) blocks for neutral stimuli only

## References

1 Frühholz, S., Trost, W. & Grandjean, D. The role of the medial temporal limbic system in processing emotions in voice and music. Progress in Neurobiology 123, 1–17, doi:10.1016/j.pneurobio.2014.09.003 (2014).

2 Pannese, A., Grandjean, D. & Fruhholz, S. Subcortical processing in auditory communication. Hear. Res. 328, 67–77, doi:10.1016/j.heares.2015.07.003 (2015).

3 Frühholz, S., Ceravolo, L. & Grandjean, D. Specific brain networks during explicit and implicit decoding of emotional prosody. Cerebral Cortex 22, 1107–1117 (2012).

4 Belyk, M., Brown, S., Lim, J. & Kotz, S. A. Convergence of semantics and emotional expression within the IFG pars orbitalis. Neuroimage 156, 240–248, doi:https://doi.org/10.1016/j.neuroimage.2017.04.020 (2017).

5 Wildgruber, D. et al. Distinct frontal regions subserve evaluation of linguistic and emotional aspects of speech intonation. Cerebral Cortex 14, 1384–1389 (2004).

6 Fruhholz, S. & Grandjean, D. Multiple subregions in superior temporal cortex are differentially sensitive to vocal expressions: a quantitative meta-analysis. Neurosci. Biobehav. Rev. 37, 24–35, doi:10.1016/j.neubiorev.2012.11.002 (2013).

7 Fruhholz, S. & Grandjean, D. Processing of emotional vocalizations in bilateral inferior frontal cortex. Neurosci Biobehav Rev 37, 2847–2855, doi:10.1016/j.neubiorev.2013.10.007 (2013).

8 Frühholz, S. & Grandjean, D. Towards a fronto-temporal neural network for the decoding of angry vocal expressions. NeuroImage 62, 1658–1666 (2012).

9 Fruhholz, S., Trost, W. & Kotz, S. A. The sound of emotions-Towards a unifying neural network perspective of affective sound processing. Neurosci. Biobehav. Rev. 68, 96–110, doi:10.1016/j.neubiorev.2016.05.002 (2016).

10 Dricu, M. & Fruhholz, S. Perceiving emotional expressions in others: Activation likelihood estimation meta-analyses of explicit evaluation, passive perception and incidental perception of emotions. Neurosci. Biobehav. Rev. 71, 810–828, doi:10.1016/j.neubiorev.2016.10.020 (2016).

11 Schirmer, A. & Kotz, S. A. Beyond the right hemisphere: Brain mechanisms mediating vocal emotional processing. Trends in Cognitive Sciences 10, 24–30 (2006).

12 Ethofer, T. et al. Cerebral pathways in processing of affective prosody: a dynamic causal modeling study. Neuroimage 30, 580–587 (2006).

13 Friederici, A. D. The cortical language circuit: from auditory perception to sentence comprehension. Trends in Cognitive Sciences 16, 262–268, doi:10.1016/j.tics.2012.04.001 (2012).

14 Kotz, S. A. et al. On the lateralization of emotional prosody: An event-related functional MR investigation. Brain and Language 86, 366–376 (2003).

15 Ethofer, T. et al. Differential influences of emotion, task, and novelty on brain regions underlying the processing of speech melody. Journal of Cognitive Neuroscience 21, 1255–1268 (2009).

16 Bach, D. R. et al. The effect of appraisal level on processing of emotional prosody in meaningless speech. NeuroImage 42, 919–927 (2008).

17 Fecteau, S., Armony, J. L., Joanette, Y. & Belin, P. Sensitivity to voice in human prefrontal cortex. Journal of Neurophysiology 94, 2251–2254 (2005).

18 Frühholz, S. & Grandjean, D. Processing of emotional vocalizations in bilateral inferior frontal cortex. Neuroscience & Biobehavioral Reviews 37, 2847–2855, doi:http://dx.doi.org/10.1016/j.neubiorev.2013.10.007 (2013).

19 Crockford, C., Gruber, T. & Zuberbühler, K. Chimpanzee quiet hoo variants differ according to context. Royal Society Open Science 5, 172066 (2018).

20 Gruber, T. & Grandjean, D. A comparative neurological approach to emotional expressions in primate vocalizations. Neuroscience & Biobehavioral Reviews 73, 182–190 (2017).

21 Beaucousin, V. et al. FMRI study of emotional speech comprehension. Cereb. Cortex 17, 339–352 (2007).

22 Mitchell, R. L. How does the brain mediate interpretation of incongruent audi-tory emotions? The neural response to prosody in the presence of conflictinglexico-semantic cues. Eur. J. Neurosci. 24, 3611–3618 (2006).

23 Fruhholz, S. et al. Neural decoding of discriminative auditory object features depends on their socio-affective valence. Soc. Cogn. Affect. Neurosci. 11, 1638–1649, doi:10.1093/scan/nsw066 (2016).

24 Villringer, A., Planck, J., Hock, C., Schleinkofer, L. & Dirnaql, U. Near infrared spectroscopy (NIRS): a new tool to study hemodynamic changes during activation of brain function in human adults. Neurosci. Lett 1, 101–104 (1993).

25 Chance, B., Zhuang, Z., Unah, C., Alter, C. & Lipton, L. Cognition-activated low-frequency modulation of light absorption in human brain. Proc Natl Acad Sci 8, 3770–3774 (1993).

26 Hoshi, Y. & Tamura, M. Detection of dynamic changes in cerebral oxygenation coupled to neuronal function during mental work in man. Neurosci Lett 1993, 5–8 (1993).

27 Kato, T., Kamei, A., Takashima, S. & Ozaki, T. Human visual cortical function during photic stimulation monitoring by means of near-infrared spectroscopy. J. Cereb Blood Flow Metab 3, 516–520 (1993).

28 Boas, D. A., Elwell, C. E., Ferrari, M. & Taga, G. Twenty years of functional near-infrared spectroscopy: Introduction for the special issue. Neuroimage 85, 1–5 (2014).

29 Homae, F. A brain of two halves: Insights into interhemispheric organization provided by Near-infrared Spectroscopy. Neuroimage 85, 354–362 (2014).

30 Buss, A. T., Fox, N., Boas, D. A. & Spencer, J. P. Probing the early development of visual working memory capacity with functional near-infrared spectroscopy. Neuroimage 85, 314–325 (2014).

31 Doi, H., Nishitani, S. & Shinohara, K. NIRS as a tool for assaying emotional function in the prefrontal cortex. Frontiers in Human Neuroscience 7, doi:10.3389/fnhum.2013.00770 (2013).

32 Bendall, R. C. A., Eachus, P. & Thompson, C. A Brief Review of Research Using Near-Infrared Spectroscopy to Measure Activation of the Prefrontal Cortex during Emotional Processing: The Importance of Experimental Design. Frontiers in Human Neuroscience 10, doi:10.3389/fnhum.2016.00529 (2016).

33 Köchel, A. et al. Affective perception and imagery: A NIRS study. International Journal of Psychophysiology 80, 192–197, doi:http://dx.doi.org/10.1016/j.ijpsycho.2011.03.006 (2011).

34 Schneider, S. et al. Show me how you walk and I tell you how you feel — a functional near-infrared spectroscopy study on emotion perception based on human gait. Neuroimage 85, 380–390 (2014).

35 Hoshi, Y. et al. Recognition of Human Emotions from Cerebral Blood Flow Changes in the Frontal Region: A Study with Event-Related Near-Infrared Spectroscopy. Journal of Neuroimaging 21, e94–e101, doi:10.1111/j.1552-6569.2009.00454.x (2011).

36 Glotzbach, E. et al. Prefrontal Brain Activation During Emotional Processing: A Functional Near Infrared Spectroscopy Study (fNIRS). The Open Neuroimaging Journal 5, 33–39, doi:10.2174/1874440001105010033 (2011).

37 Tupak, S. V. et al. Implicit emotion regulation in the presence of threat: Neural and autonomic correlates. Neuroimage 85, 372–379 (2014).

38 Herrmann, M. J., Ehlis, A. C. & Fallgatter, A. J. Prefrontal activation through task requirements of emotional induction measured with NIRS. Biol. Psychol. 64, 255–263 (2003).

39 Aoki, R. et al. Relationship of negative mood with prefrontal cortex activity during working memory tasks: An optical topography study. Neuroscience Research 70, 189–196, doi:http://dx.doi.org/10.1016/j.neures.2011.02.011 (2011).

40 Ozawa, S., Matsuda, G. & Hiraki, K. Negative emotion modulates prefrontal cortex activity during a working memory task: a NIRS study. Frontiers in Human Neuroscience 8, doi:10.3389/fnhum.2014.00046 (2014).

41 Plichta, M. M. et al. Auditory cortex activation is modulated by emotion: A functional near-infrared spectroscopy (fNIRS) study. Neuroimage 55, 1200–1207, doi:http://dx.doi.org/10.1016/j.neuroimage.2011.01.011 (2011).

42 Zhang, D., Zhou, Y. & Yuan, J. Speech prosodies of different emotional categories activate different brain regions in adult cortex: an fNIRS study. Scientific Reports 8, 218, doi:10.1038/s41598-017-18683-2 (2018).

43 Tsuzuki, D. et al. Virtual spatial registration of stand-alone fNIRS data to MNI space. NeuroImage 34, 1506–1518, doi:https://doi.org/10.1016/j.neuroimage.2006.10.043 (2007).

44 Frühholz, S., Klaas, H., Patel, S. & Grandjean, D. Talking in fury: the cortico-subcortical network underlying angry vocalizations. Cereb. Cortex. 25, 2752–2762 (2015).

45 Jasper, H. H. The Ten-Twenty Electrode System of the International Federation. Electroencephalography and Clinical Neurophysiology 10, 371–375 (1958).

46 Okamoto, M. et al. Three-dimensional probabilistic anatomical cranio-cerebral correlation via the international 10–20 system oriented for transcranial functional brain mapping. NeuroImage 21, 99–111, doi:https://doi.org/10.1016/j.neuroimage.2003.08.026 (2004).

47 Delpy, D. T. et al. Estimation of optical pathlength through tissue from direct time of flight measurement. Phys Med Biol 33, 1433–1442 (1988).

48 Scholkmann, F., Spichtig, S., Muehlemann, T. & Wolf, M. How to detect and reduce movement artifacts in near-infrared imaging using moving standard deviation and spline interpolation. Physiol. Meas. 31, 649–662 (2010).

49 Oostenveld, R., Fries, P., Maris, E. & Schoffelen, J.-M. FieldTrip: Open Source Software for Advanced Analysis of MEG, EEG, and Invasive Electrophysiological Data. Computational Intelligence and Neuroscience 2011, 9, doi:10.1155/2011/156869 (2011).

50 Friston, K. J., Mechelli, A., Turner, R. & Price, C. J. Nonlinear responses in fMRI: The balloon model, volterra kernels, and other hemodynamics. Neuroimage 12, 466–477 (2000).

51 Ethofer, T. et al. Cerebral pathways in processing of affective prosody: a dynamic causal modeling study. Neuroimage 30, 580–587, doi:S1053-8119(05)00728-7[pii] 10.1016/j.neuroimage.2005.09.059 (2006).

